# Identification and functional prediction of sugar beet circRNAs involved in drought responses

**DOI:** 10.1101/2022.08.03.502711

**Authors:** Chun-Lei Zou, Zhiqiang Guo, Shanshan Zhao, Jishuai Chen, Chunlai Zhang

## Abstract

Drought is one of the most common abiotic constraints on the quality and productivity of crops on a global scale. Despite the rapidly updating information on circRNAs (circular RNAs), their roles in the anti-drought regulation of sugar beet are least understood. As a newly recognized class of non-coding RNAs, circRNAs exert crucial effects on miRNA (microRNA) functionality, as well as on transcriptional regulation. To clarify the mechanism of how circRNAs of sugar beet respond to drought stress, deep sequencing was employed to characterize these circRNAs in a genome-wide manner under drought treatment. Our results identify a total of 17 differentially expressed circRNAs. As revealed by the Kyoto Encyclopedia of Genes and Genomes and Gene Ontology outcomes, circRNAs were found capable and involved in drought-responsive events. Utilizing the target genes exhibiting direct/indirect associations with drought resistance, we established a circRNA–miRNA–mRNA meshwork based on the circRNAs that were expressed differentially. The probable sponge functions of *novel_circ_0000442* and *novel_circ_0000443* were exerted by targeting *ath-miR157d*. This helped regulate the expression of relevant target genes, including *BVRB_1 g004570, BVRB_1 g005450*, and *BVRB_1 g005790*, that were involved in drought response. Apart from offering novel understandings of anti-drought mechanisms, our findings lay a basis for probing deeper into the intricate regulatory networks of sugar beet genes.

## Introduction

Drought is the most common abiotic constraint on the quality and productivity of crops. The severity and frequency of drought have been increasing globally in recent decades because of greenhouse effect-induced climate change. Drought stress in plants can trigger detrimental reactions, including membrane system disruption, osmotic imbalance, as well as declined photosynthetic and respiratory rates. These affect plant growth and metabolism throughout the growth stages, which compromise crop productivity and quality (Liu *et al*., 2020b). Plants respond intricately to drought stress, which involves not only the physiological and cellular dimensions but also the molecular dimension (Giordano *et al*., 2016). Similar to the response to other abiotic stresses, multi-gene interactions via various pathways are required for initiating a response to drought stress at the molecular level. Plants differentially express a few regulatory and functional genes under drought stress. This establishes an intricate meshwork of signal regulation which influences a range of their biochemical and physiological responses (Wang *et al*., 2021a). For instance, water deficiency leads to restriction in photosynthesis because of damage to the photosynthetic machinery, leaf expansion lessening, as well as reduced activities of Calvin cycle enzymes like phosphoenolpyruvate and Rubisco carboxylases (Bota *et al*., 2004). ROSs (reactive oxygen species) generated in a water deficiency context lead to unstable and senescent membranes of cells or the death of plants by targeting a multitude of organelles like chloroplasts, peroxisomes, and mitochondria (Ma *et al*., 2013). Hormonal regulation is another crucial factor enabling water deficit tolerance in plants. To diminish the detrimental impacts resulting from water deficiency, expressions of a few transcriptional factors are induced. At the same time, their target genes are implicated in the activation of ABA (phytohormone abscisic acid), as well as critical signaling and perception components (Ahuja *et al*., 2010; Sirichandra *et al*., 2009).

A novel and potent technique for unraveling the phenotype-genotype association is high-throughput sequencing. RNA-seq, for instance, has gained extensive usage in plant genetics. Transcriptome analysis, in particular, is applied for unraveling DEGs (differential expression genes) in diverse biological events (Hong *et al*., 2020). Accompanying the progression of high-throughput sequencing and highly-effective big data analysis, a growing number of ncRNAs (non-coding RNAs) has been recognized and elucidated in plants under stress (Zhang, 2015). Depending upon the length, they can be classified into siRNAs (small interfering RNAs), circRNAs (circular RNAs), lncRNAs (long non-coding RNAs), as well as small RNAs like miRNAs (micro RNAs) (Liu *et al*., 2022b; Wang *et al*., 2020; Zou *et al*., 2021). Through direct interplays with RNAs, DNAs, and proteins, the non-coding RNAs are capable of regulating the expression of the genes responsive to environmental stresses (Khaldun *et al*., 2016).

The circRNAs, which are a special kind of non-coding RNAs without 3’tails or 5’caps, have been identified that accompanied progression in high-throughput sequencing and highly-effective big data analysis. (Chen *et al*., 2018). Innumerable circRNAs have been recognized both in humans and animals (Legnini *et al*., 2017; Memczak *et al*., 2013). Lately, circRNAs have been widely studied in plants like Arabidopsis (*A. thaliana*) (Chen *et al*., 2017; Liu *et al*., 2017), rice (*Oryza sativa*) (Ye *et al*., 2015), tomato (*Lycopersicon esculentum*) (Yin *et al*., 2018), wheat (*Triticum aestivum*) (Wang *et al*., 2017), and soybean (*G. max*) (Zhao *et al*., 2017). The probable implication of circRNAs in the chilling responsive event of tomato has been reported (Zuo *et al*., 2016), while cold tolerance-associated circRNA has been identified in grapes(Gao *et al*., 2019). These imply the cold stress-regulatory actions of plant circRNAs.

A crucial commercial crop called *Beta vulgaris L. (sugar beet*) contributes greatly to the worldwide supply of sugar. Traits particularly linked to its productivity enhancement are adaptations to both biotic and abiotic stresses, including cold in a temperate environment, drought, heat, as well as salinity (Taleghani *et al*., 2022; Zou *et al*., 2020). Drought adaptation refers to a process in which the sugar beet develops fundamental resistance against drought challenges through biochemical and physiological alterations (Alkahtani *et al*., 2021; Li *et al*., 2019). The drought response of sugar beet is definitively linked to the levels of *BvHb2* and other protein-encoding genes (Gisbert *et al*., 2020). Nevertheless, there is insufficient knowledge about the effects of non-coding RNAs in drought response of sugar beet. In the present work, circRNAs in leaves of sugar beet were assessed by employing high-throughput sequencing combined with bioinformatic measures in a water control context to investigate the genome-wide quantity of circRNAs, as well as their possible drought response-regulatory effects. For this role, a circRNAs-miRNAs–mRNAs meshwork was created, while for the expression pattern exploration of drought-associated sugar beet circRNAs, quantitative real-time PCR was employed. Our findings will be of potential significance to the complexity evaluation of the regulatory circRNAs concerning the plant responses to drought.

## Materials and methods

### Plant materials and treatment

Following pot germination, the seeds of sugar beet variety KWS9147 were subjected to cultivation under 25 °C and 80% RH conditions, where the photoperiod was 16 h (day)/8 h (night). At the phase of full leaf expansion, the seedlings were assigned to either (1) well-watered (CK) or (2) drought stress (DR) group. Water was added to the control group up to 70% of the holding capacity while maintaining normal growth, whereas, in the drought group, watering was prohibited. Three independent biological replicates (10 plants for each replicate) were performed.On the 8th day following the drought challenge, when the pronounced curling of the leaves was noted, sample harvesting was accomplished. From every treatment group, leaf cuttings were acquired, placed immediately into the liquid nitrogen, and then subjected to a –80 °C cryopreservation.

### Determination of proline content, catalase activity, and abscisic acid content in leaves

The sulfosalicylic acid approach was employed for assessing proline content (Marques de Carvalho *et al*., 2021). Assaying of catalase (CAT) activity was accomplished as per Liu *et al*.’s procedure (2021). One unit of CAT activity referred to the quantity of enzyme needed for decomposing 1 μmol of H_2_O_2_ min^−1^ mL^−1^.

Abscisic acid (ABA) was extracted and quantified from sugar beet leaves by a modified version of Brunetti *et al*.’s procedure (2020). Initially, following liquid nitrogen-based pulverization and mixing with d6-ABA (50 ng), the leaves (0.5 g) were subjected to a 30-min extraction using CH_3_OH/H_2_O (50:50; v:v; pH 2.5; 3 × 1 mL) at 4 °C. Next, the supernatant was subjected sequentially to n-hexane (3 × 3 mL) defatting, the Sep-Pak C18 cartridge (Waters, Milford, USA) purification, and ethyl acetate (1 mL) elution. After drying under nitrogen, the eluate was rinsed using CH_3_OH/H_2_O (50:50; pH 2.5; 500 μL) and subsequently loaded (3 μL aliquots) onto an LC–DAD-MS/MS system comprising an LCMS-8030 quadrupole MS and a Nexera HPLC (both Shimadzu, Kyoto, Japan), whose operational mode was ESI (electrospray ionization). The eluting phases comprised H_2_O involving HCOOH (0.1%) plus solvent A and CH_3_CN/CH_3_OH (1:1, v:v) involving HCOOH (0.1%) plus solvent B. Under the negative ion conditions, a Poroshell 120 SB C18 column (3 × 100 mm, 2.7 μm, 4.6 × 100 mm, Agilent, Palo Alto, USA) was utilized during the analysis. An 18-min elution from solvent A (95%) to solvent B (100%) was accomplished at a 0.3 mL min^−1^ flow rate. The MRM (multiple reaction mode) was adopted for the quantitative assessment.

### RNA extraction, quantification, and qualification

Extraction of total RNA was accomplished as per the protocol of TRIzol reagent (Invitrogen, Carlsbad, USA). For the elimination of genomic DNA contaminants, DNase I (Takara Bio, Dalian, China) was used to process the total RNA, followed by a 1-hour incubation of purified DNase I-processed total RNA. It was then proceeded using RNase R (3 U/μg; Epicentre, Madison, USA) at 37 °C. Finally, with the aid of Bioanalyzer 2100 system (Agilent, CA, USA), overall RNA quantity and integrity were evaluated via the RNA Nano 6000 Assay Kit.

### Library preparation and circRNA sequencing

RNA libraries were constructed using ribosomal-depleted RNAs as per the protocol of Ultra Directional RNA Library Prep Kit for Illumina (NEBNext, NEB, USA). The random hexamer primer plus RNaseH-(M-MuLV Reverse Transcriptase) was utilized for the synthesis of strand cDNA. For succeeding synthesis of 2nd strand cDNA, RNase H combined with DNA Polymerase I was utilized. Following the preferential selection of 250–300-bp-long fragments, the PCR (polymerase chain reaction) amplification proceeded for the yielded samples.

Through trimming of ploy-N-or adapter-containing reads or the low-quality reads from raw data, the acquisition of clean data (reads) was accomplished. Meanwhile, the GC, Q20, and Q30 contents were estimated for the clean data. The high-quality clean data were utilized to make the entire downstream analyses. Annotation files of the gene model and reference genome were downloaded from the genome website. Then, Bowtie was exploited to align the high-quality clean reads to the reference genome (*Beta_vulgaris_Ensembl*) (Langmead *et al*., 2009). Examination and recognition of circRNA were accomplished via find_circ (Memczak *et al*., 2013) and CIRI2(Gao *et al*., 2017). For circRNA expression assessment, the reads per million mapped (RPM) was adopted.

### Analysis of differential expression of circular RNAs

With the aid of DESeq R ver. 1.24.0, differential expression evaluation was accomplished between two groups or conditions (Wang *et al*., 2010). The circRNA levels from triplicate experiments were averaged to serve as the outcome of one treatment. The paired t-test was employed to recognize the circRNAs that were expressed significantly differentially between the groups, where the threshold was |log_2_FC| (fold change/FC) > 1 and the p-values were < 0.05.

QRT-PCR (quantitative real-time PCR) assessment was performed on 12 differentially expressed circRNAs under CK and DR conditions, intending to verify the expression patterns of RNA-seq-identified circRNAs. The back splice junction was crossed through the primer (Premier 5.0-based design of circRNA primers) so that the circular templates’ head-to-tail junctions could be amplified (Shen *et al*., 2015). Supplementary Table S1 details the sequences for the entire sequences. The qRT-PCR detection system (LineGene 9600 Plus, BIOER., Hangzhou, China) was utilized for qRT-PCR assays, where the miRcute SYBR Green MasterMix and SYBR GreenMaster Mix (both Tiangen, Beijing, China) were used. Computation of data was achieved by 2 approach. Data were normalized to the Actin, the housekeeping gene for the sugar beet. Triplicate assays of qRT-PCR were performed, and the outcomes were represented in terms of mean ±SE.

### GO and KEGG enrichment analysis

For potential functionality assessment of circRNAs’ parental genes, KEGG (Kyoto Encyclopedia of Genes and Genomes) and the GO (Gene Ontology) databases were utilized for annotating the parental genes. GOseq R was exploited to make GO enrichment assessment on the parental genes of circRNAs that expressed differentially, where correction of length bias was implemented for the genes (Young *et al*., 2010). GO terms were regarded as significantly enriched by differential expressed genes when corrected p-values were < 0.05. There were three ontologies for the GO database: molecular functionality, biological event, and the cellular component. During the pathway enrichment assay, metabolic or signaling pathways in the parental genes that were pronouncedly enriched in contrast to the entire genome background were identified. The pathways in parental genes were deemed significantly enriched when p-values were < 0.05.

### CircRNA-miRNA-gene network analysis

The miRanda (animal species) or target finder (plant species) was exploited for identifying the MicroRNA target sites in exons of circRNA loci. The meshwork of circRNA–miRNA-gene was established with the aid of Cytoscape.

## Results

### The effect of drought stress on some physiological traits

To examine how drought stress influenced the physiological traits of our test plant, proline, ABA levels, as well as CAT activity were determined in the leaves acquired from plants under CK and DR treatments. It is apparent (Fig. 1) that these three physiological traits were elevated prominently by the drought stress. As implied by the aforementioned physiological heterogeneities, the expressions of sugar beet genes, including circRNAs, altered following drought stress.

**Fig. 1.**
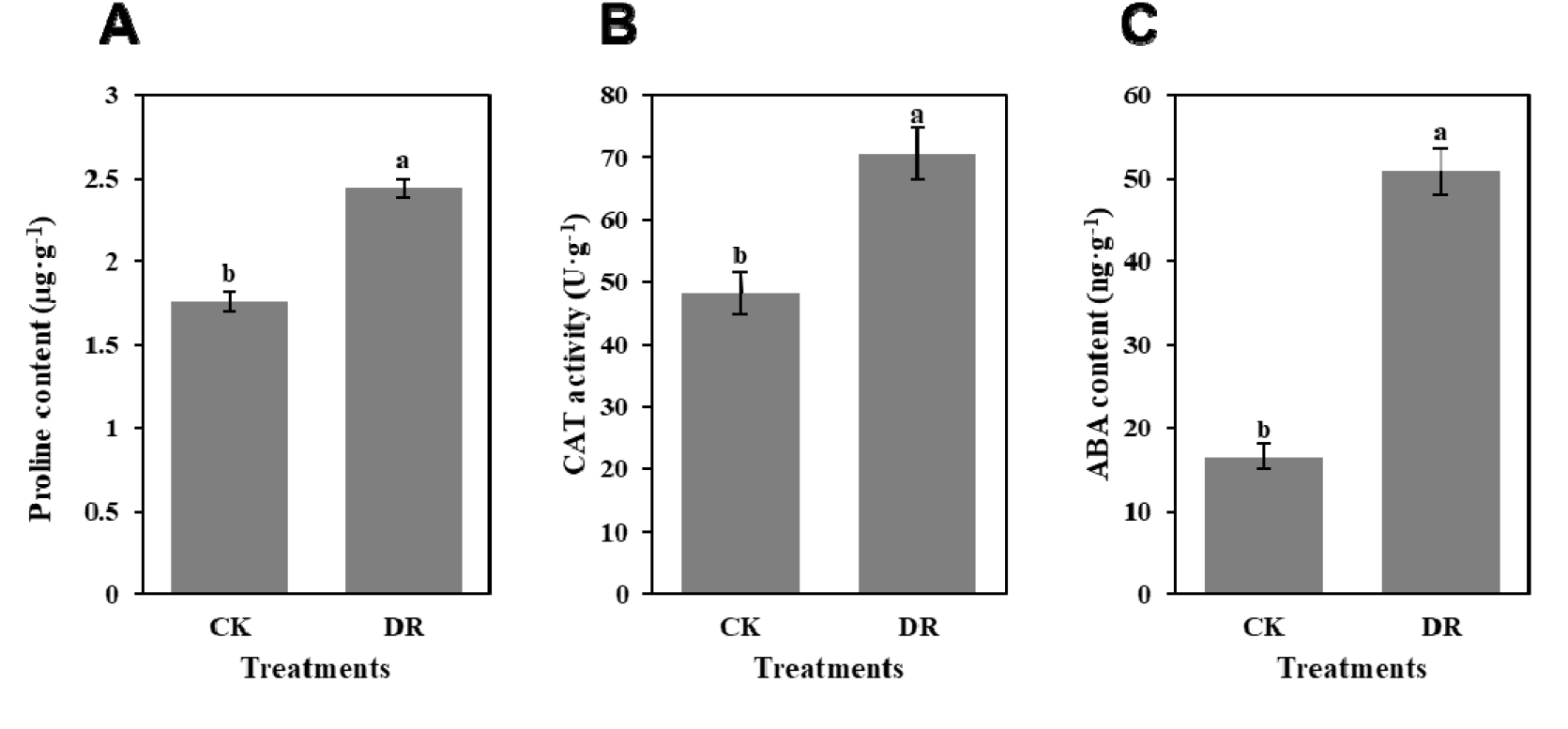
Effect of drought stress on some physiological traits of sugar beet. (A) Proline content. (B) CAT activity. (C) ABA content. Means ±SE are presented.

### Identification of circRNAs in sugar beet

We created and sequenced a total of six RNA-seq libraries (Table 1) for the present study. The number of novel circRNAs identified from our circRNA-seq data totaled 565. Supplementary Table S2 presents the details of the circRNAs identified in this study.

**Table 1.**
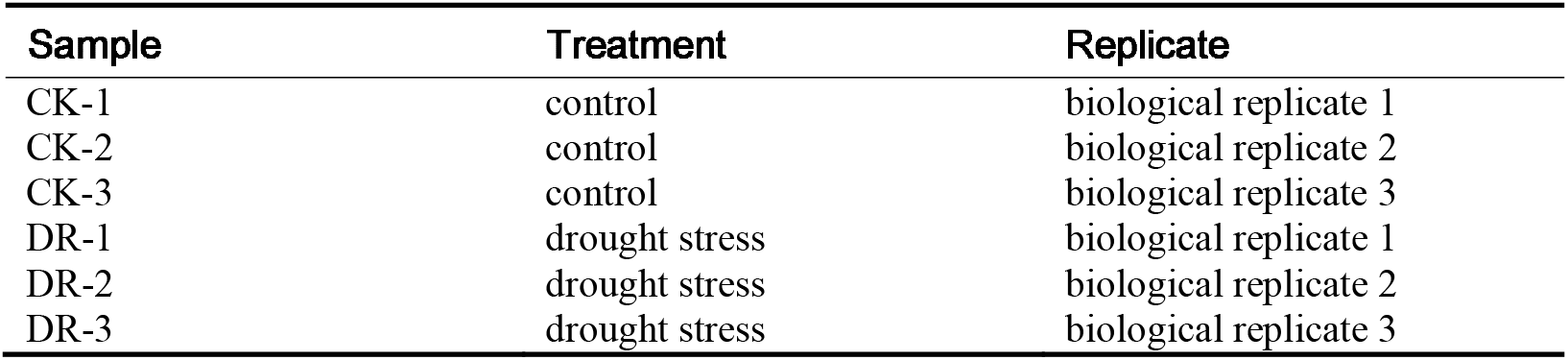
Profile of sample information.

The 563 circRNAs were categorized into three types, viz. exon, intron, and intergenic depending on their genomic origin.: The total number of the exon, intron, and intergenic types of circRNAs were 432, 17, and 114, respectively (Fig. 2A). The exon-type circRNAs were the prevailing type, representing about 76.7% of the entire 563 circRNAs. These outcomes agree with the findings in other species like *Oryza sativa, Brassica rapa*, and *Arabidopsis thaliana*, whose circRNAs originated from the protein-encoding genes’ exons, separately representing, respectively, 85.7%, 52.0%, and 50.5% of the total (Liu *et al*., 2022a; Weber *et al*., 1999). As displayed in Fig. 2B, the distribution range of the number of these circRNAs on the sugar beet chromosomes was 36–70.

**Fig. 2.**
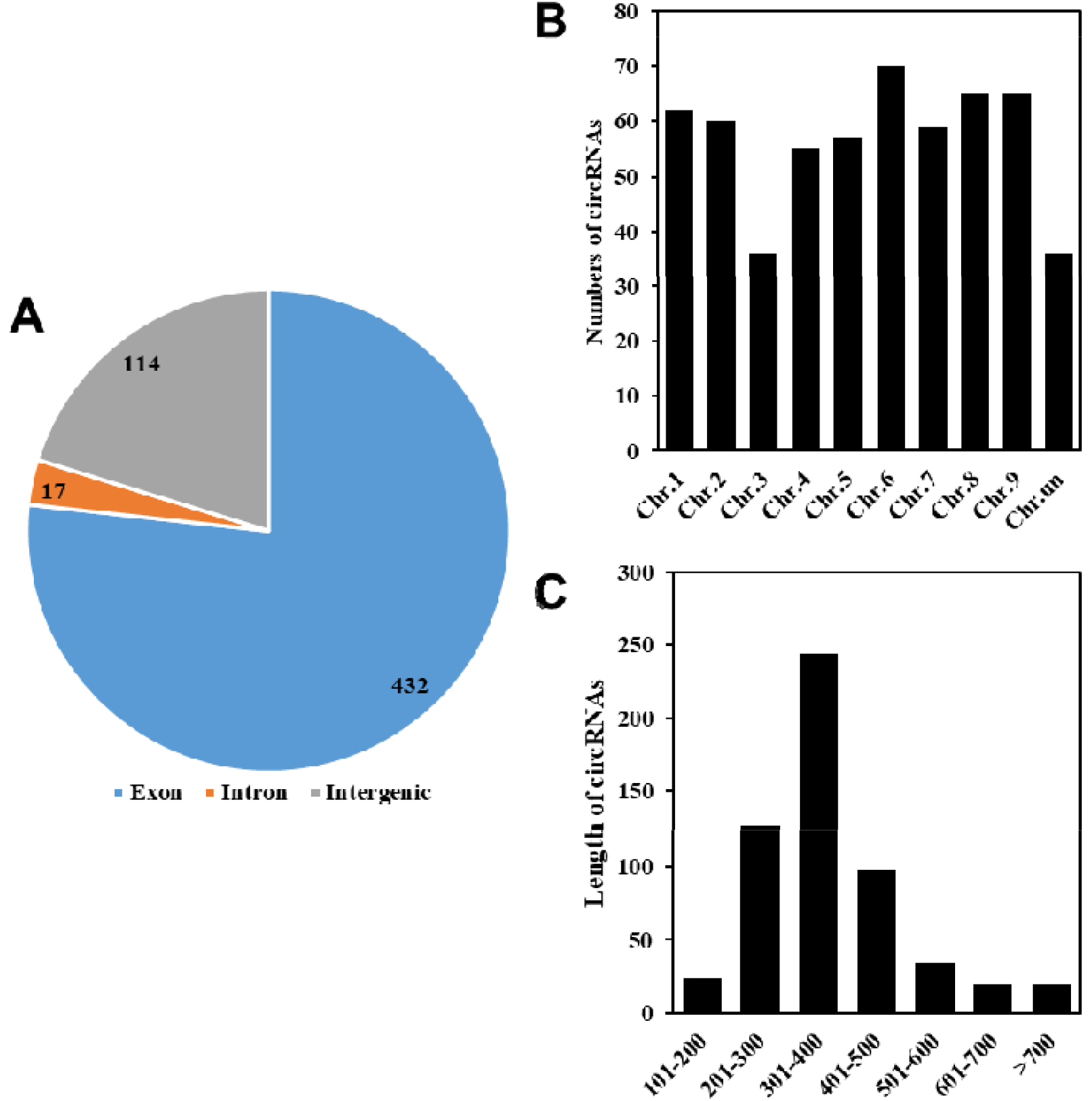
Characterization of sugar beet circRNAs. (A) Types of circRNAs. (B) Histogram of the distribution of circRNAs on the chromosomes. (C) Distribution of the length of the circRNAs.

For instance, 70 circRNAs from Chr.6 were the most predominant ones (12.4%), followed by Chr.8 and Chr.9. For these circRNAs, the length range was 101–700 bp. Moreover, the majority of them were 301–400 bp in length (Fig. 2C).

### Identification of CircRNA Parental Genes

We defined the annotated circRNA-generating genes as the circRNAs’ parental genes, while “NA” represented the circRNAs having no parental genes. According to Supplementary Table S2, the origin of 527 circRNAs was associated with 448 parental genes, while 36 intergenic circRNAs originated from the fragments between two genes and therefore had no specific parental genes. The number of circRNAs originating from a single parental gene totaled 526. There was only one circRNA (*novel_circ_0000910*) with more than one parental gene. Given the alternative splicing pattern of sugar beet, varying circRNAs were generated by the parental genes. This finding coincided with the previous works indicating alternative splicing patterns possessed by circRNAs. These are thus a valuable resource for comprehending the intricate biogenesis of circRNAs, as well as their potential functionalities (Gao *et al*., 2016; Zhang *et al*., 2016).

### Differential circRNA expression patterns of sugar beet in response to drought stress

As displayed in Fig. 3A, among the 563circRNAs, 518 were identified in the CK group (including 348 specifically expressed unique ones), whereas 215 were identified in the DR group (including 45 specifically expressed unique ones). The number of significantly differentially expressed (DE) circRNAs between these 2 groups totaled 17 (Fig. 3B). These specific expression profiles offered clues about the bio-functionalities of circRNAs. For the exploration of the expression of circRNAs from sugar beet in the DR scenario, qRT-PCR assays were performed on 12 randomly chosen circRNAs, as presented in detail in Supplementary Table S1. The expression patterns of these 12 circRNAs agreed with the outcomes of RNA-seq, of which nine were upregulated, and three were down-regulated (Fig. 3B; Fig. 4). Over 2-fold differences were noted in the expression levels of several circRNAs, such as *novel_circ_0000853*, novel_circ_0000695, *novel_circ_0000043 andnovel_circ_0000112*; and an over 10-fold elevation was noted in the *novel_circ_0000591* expression levels (Fig. 4). The 12 circRNAs under drought challenge differed significantly from the control ones, suggesting probable crucial effects of these circRNAs on drought tolerance. This is implied by the pronouncedly positive linkages of the expression levels of *novel_circ_0000591*, *novel_circ_0000736, novel_circ_0000695*, and *novel_circ_0000764* to those of their respective parent genes *BVRB_7 g173230, BVRB_9 g204220, BVRB_8 g185430* and *BVRB_9 g217640* under drought conditions (Fig. 4). Therefore, circRNAs probably exerted a pivotal effect on the drought-responsive control via the *Cis*-regulation of their parent genes.

**Fig. 3.**
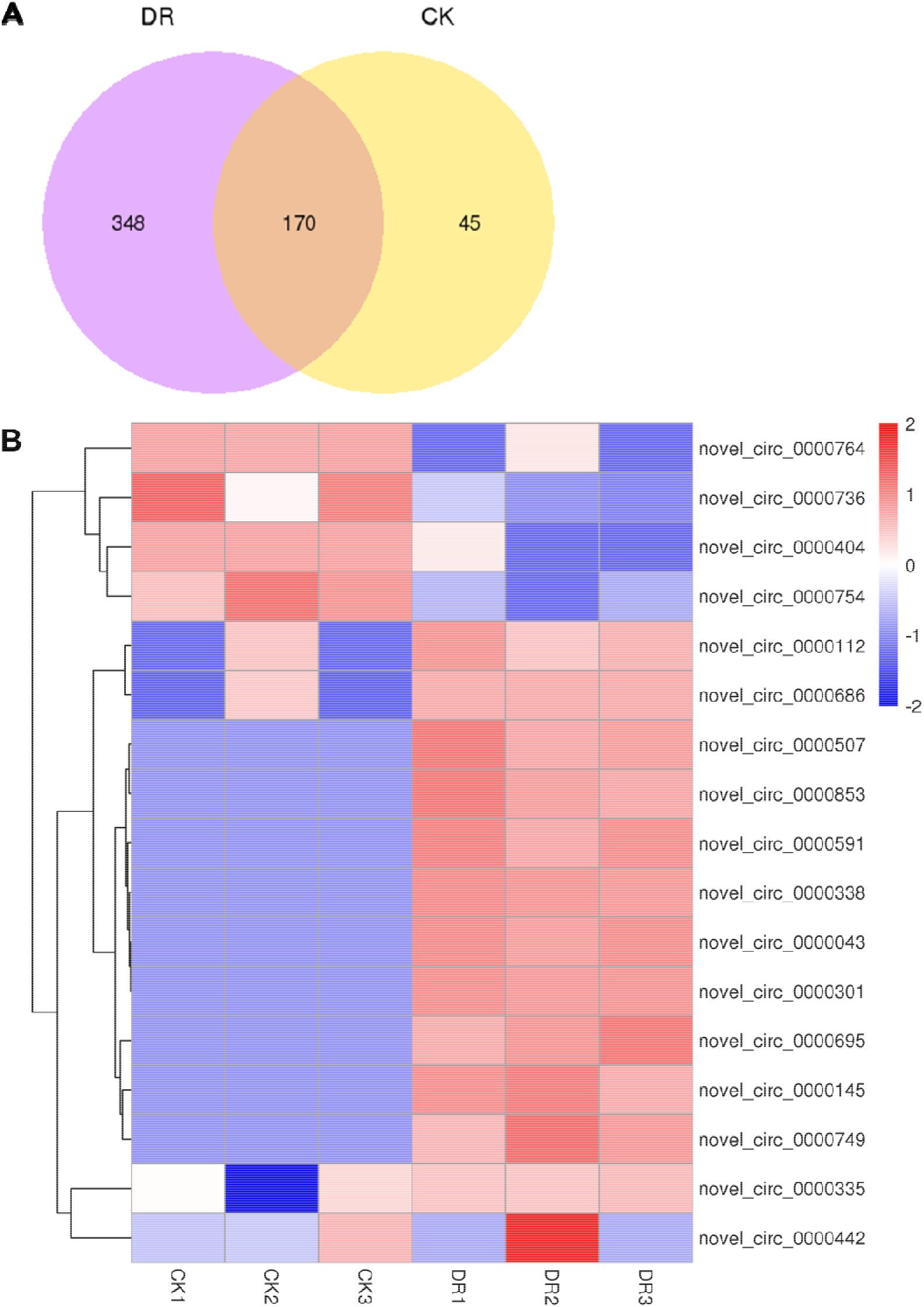
The DE circRNAs of sugar beet upon drought challenge. (A) A Venn plot describing common and specific circRNAs under DR and CK conditions. (B) A heat map illustration describing the levels of drought-responsive circRNA expression.

**Fig. 4.**
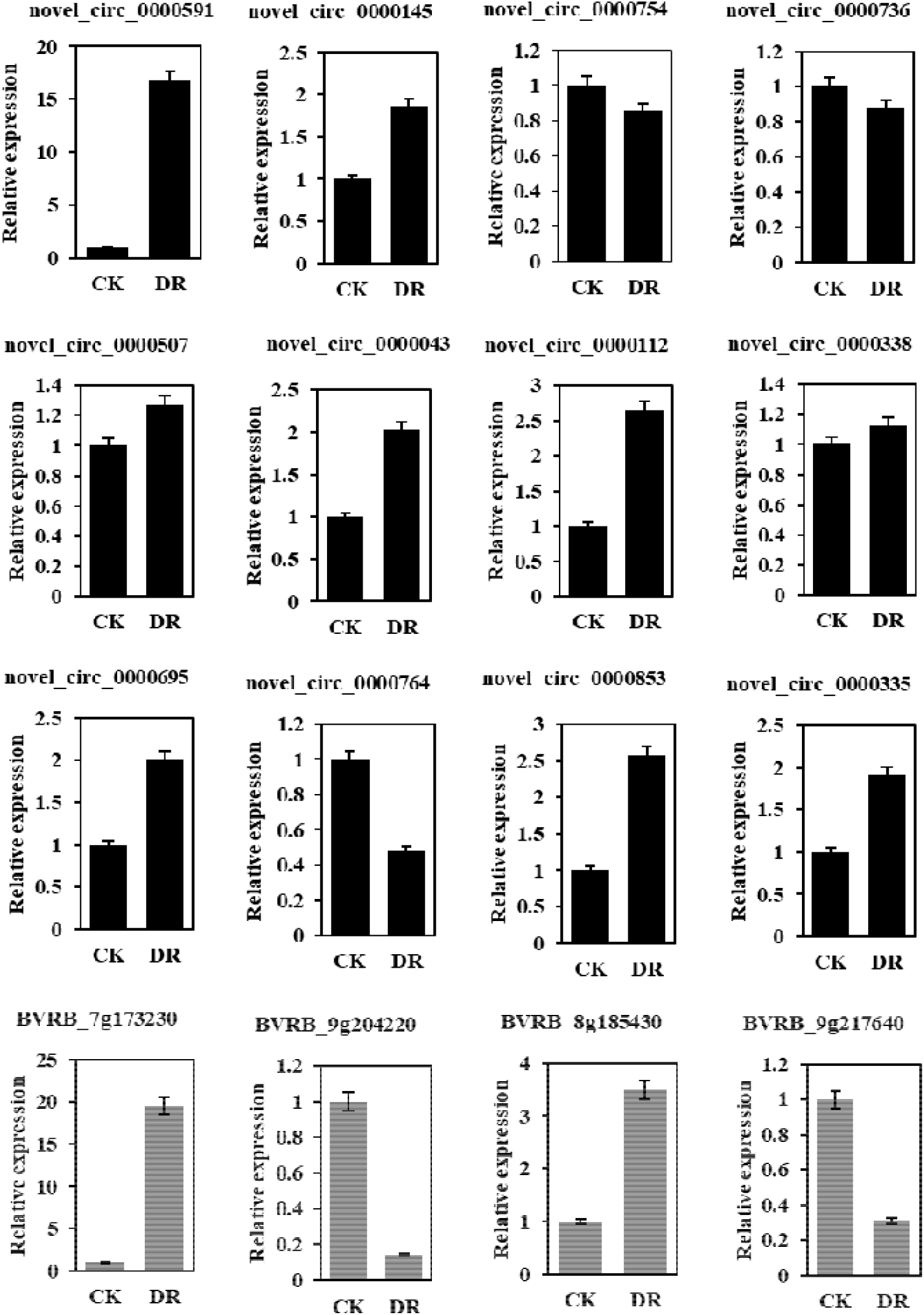
QRT-PCR-based verification of DE circRNAs and part of parental genes indicating means ±SEs from independent biological triplicate, as well as three technical repeats.

### Functional annotation for parental genes of sugar beet circRNAs in the DR scenario

The crucial effects of circRNAs on transcriptional control were reported, which were achieved through the Cis-regulation of their parental genes (Li *et al*., 2017b). These results are in agreement with the present study (Fig. 4). The parental genes of circRNAs identified herein were subjected to KEGG and GO analyses, to investigate the presumed functionality of sugar beet circRNAs under DR challenges. In biological event terms, the two most pronouncedly enriched categories were the cellular nitrogen compound metabolic process (GO:0034641) and the organic cyclic compound metabolic process (GO:1901360). Concerning the cellular components, primary implications of circRNA parental genes were noted in the transferase complex (GO: 1990234). For molecular function, nucleic acid binding (GO:0003676) was the most enriched GO term (Fig. 5A). As revealed by the KEGG assessment, there were 13 pathways linked to the drought tolerance in sugar beet, such as metabolic process-associated ones (glyoxylate, dicarboxylate, glycine, serine, threonine, fructose, mannose, cysteine, methionine, carbon, and relevant secondary metabolites), as well as those associated with plant hormonal signaling, plant-pathogen interaction, ABC transporters, glycolysis/gluconeogenesis, and pentose phosphate pathway (Fig. 5B).

**Fig. 5.**
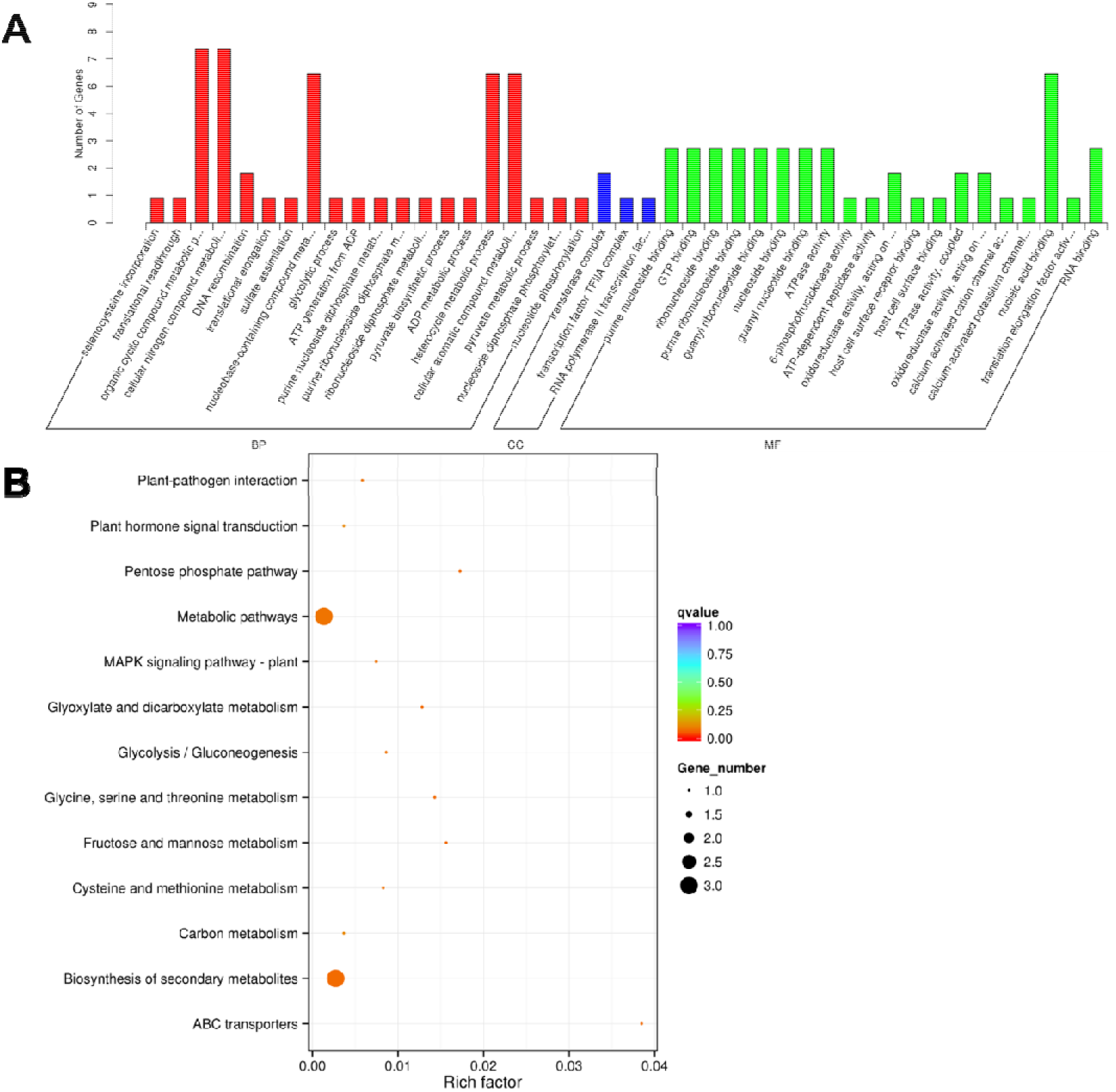
(A) The GO categorization and (B) KEGG enrichment outcomes for the parental genes of DE circRNAs upon drought challenge.

Signal transduction and oxidation-reduction play pivotal roles in plant stress response. Parent genes of some drought-responsive circRNAs were related to signal transduction. For instance, the parent gene of *novel_circ_0000591* (upregulated more than 65-fold under DR treatment) was *BVRB_7 g173230* (probable protein phosphatase 2C 24) (Table 2). There were also some parent genes involved in oxidation-reduction in the present study. For example, *BVRB_1 g013970* (member 1 of short-chain dehydrogenase/reductase family 42E), *BVRB_8 g185430* (pyrophosphate: β subunit of fructose 6-phosphate 1-phosphotransferase) and *BVRB_9 g213980* (glycerate dehydrogenase HPR, peroxisomal) were the parent genes of *novel_circ_0000112* (up-regulated more than 7-fold under DR treatment), *novel_circ_0000695* (up-regulated more than 25-fold) and *novel_circ_0000749* (up-regulated more than 26-fold). As suggested by these outcomes, drought stress is prominently influential to the redox and signal transduction in sugar beet. Furthermore, the expression level of *novel_circ_0000043*, whose parent gene was *BVRB_007520* (probable DNA helicase MCM8), was upregulated more than 29-fold under DR treatment. This phenomenon suggests that sugar beet might accelerate its DNA replication toward drought response.

**Table 2.** Parent genes of some DE circRNAs involved in stress response.

| CircRNAs ID | log2FoldChange | P-value | Host gene ID | Host gene description |
| --- | --- | --- | --- | --- |
| *novel_circ_0000591* | 6.0421 | 0.000304 | *BVRB_7 g173230* | probable protein phosphatase 2C 24 |
| *novel_circ_0000043* | 4.8777 | 0.018551 | *BVRB_007520* | probable DNA helicase MCM8 |
| *novel_circ_0000112* | 2.9851 | 0.034708 | *BVRB_1 g013970* | short-chain dehydrogenase/reductase family 42E member 1 |
| *novel_circ_0000695* | 4.679 | 0.033082 | *BVRB_8 g185430* | pyrophosphate--fructose 6-phosphate 1-phosphotransferase subunit beta |
| *novel_circ_0000749* | 4.7417 | 0.0329 | *BVRB_9 g213980* | glycerate dehydrogenase HPR, peroxisomal |

### Construction of circRNA-miRNA-mRNA networks

Recently, the crucial effects of circRNAs on downstream target gene regulation have been reported, which is achieved by sponging miRNAs and controlling gene expression (Liu *et al*., 2022a; Wang *et al*., 2020). The possible target sites of miRNAs were identified in the sugar beet circRNAs by the bioinformatic means to clarify whether the post-transcriptional levels of target genes were affected by these circRNAs via the miRNA binding. As displayed in Supplementary Table S3, we found the presence of putative miRNA-binding sites in 197 of 563 (35.0%) circRNAs, and there were 166 speculated miRNAs combined with these 197 circRNAs. Fifty-five of these 197 circRNAs possessed over one miRNA-binding site. The *novel_circ_0000196* possessed the largest quantity (14) of miRNA-binding sites, while *ath-miR414* had the largest number of circRNAs (15). Fig. 6A depicts the miRNA-binding sites of *novel_circ_0000112* and *novel_circ_0000695*, whereas Supplementary Table S3 describes their details. As suggested by these outcomes, there may be plentiful miRNA-binding sites in the circRNAs of sugar beet, which may impact the anti-drought gene expression via the miRNAs. With the aid of Cytoscape, a holistic meshwork of circRNA–miRNA–mRNA interactions was created, as illustrated in Fig. 6B.

**Fig. 6.**
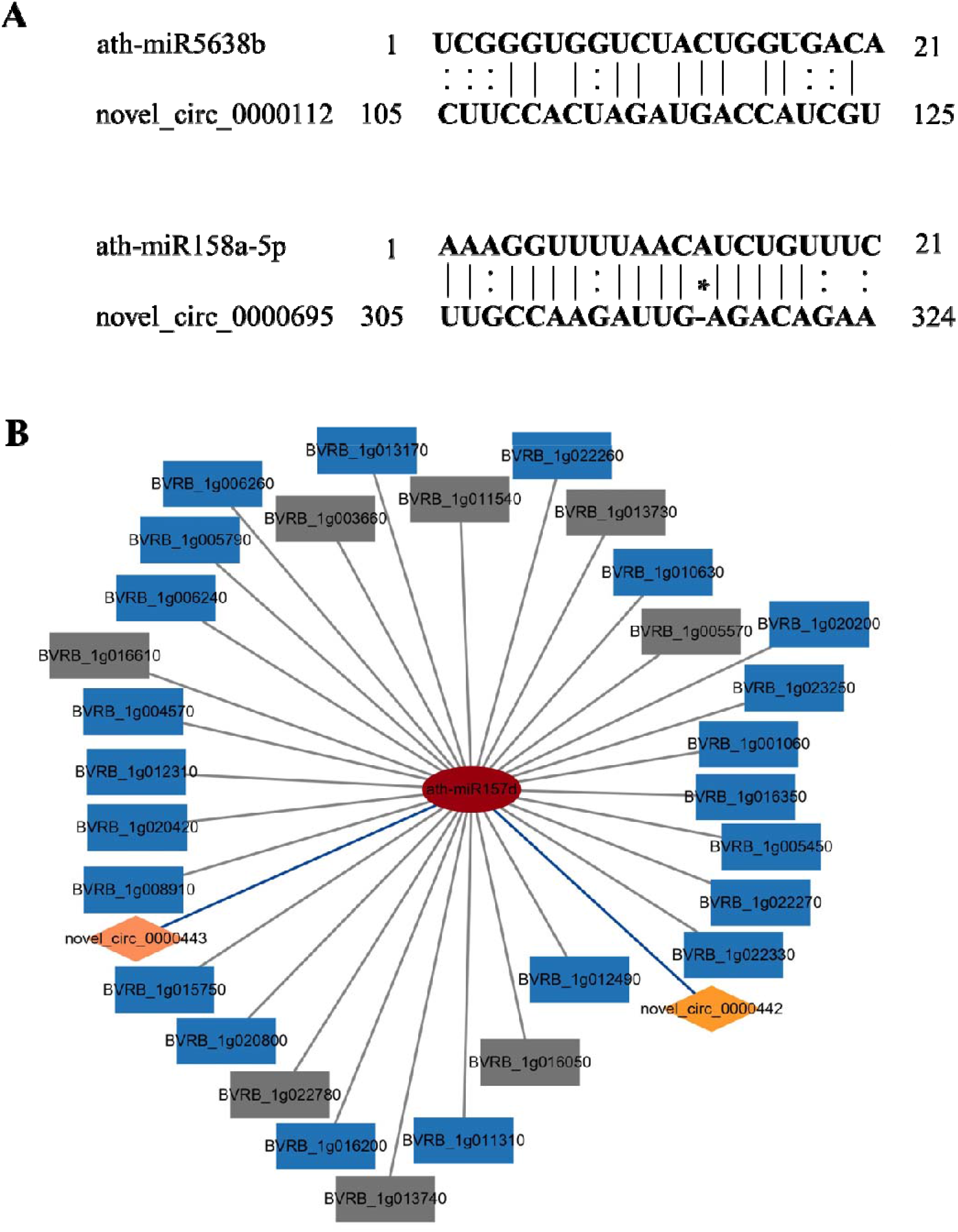
(A) Complementary base pairing diagram for the interactions between miRNAs and circRNAs. “|” stands for the AU–GC matching bases, “:” represents GU–AC matching bases, and “*” stands for the mismatching bases. (B) A meshwork of circRNA–miRNA–mRNA interplays for the sugar beet circRNAs upon drought challenge. Rhombic, rectangular and circular nodes stand separately for circRNAs, target genes, and miRNAs. Drought-responsive genes are marked by blue rectangular nodes, while other genes are marked by gray rectangular nodes.

Clearly, diverse circRNAs could target an individual miRNA. The *ath-miR157d*, for instance, was speculated to be targeted by *novel_circ_0000442* and *novel_circ_0000443*, two circRNAs that are capable of targeting a few drought-responsive genes like the U-box domain-containing protein *33BVRB_1 g004570*, protein *CLT2BVRB_1 g005450and* methionine adenosyltransferase 2 subunit beta *BVRB_1 g005790* (Fig. 6B; Table 3). Therefore, the leaf morphogenesis-related miRNA sponges of sugar beet probably exerted a crucial effect on enhancing drought resistance control.

**Table 3.** Functional description of targeted genes for *ath-miR157d* in response to drought stress.

| Target gene ID | Target description |
| --- | --- |
| *BVRB_1g001060* | Uncharacterized membrane protein At4g09580 |
| *BVRB_1g004570* | U-box domain-containing protein 33 |
| *BVRB_1g005450* | Protein CLT2 |
| *BVRB_1g005790* | Methionine adenosyltransferase 2 subunit beta |
| *BVRB_1g006240* | B-box zinc finger protein 21 |
| *BVRB_1g006260* | Probably inactive leucine-rich repeat receptor-like protein kinase IMK2 |
| *BVRB_1g008910* | Putative pentatricopeptide repeat-containing protein At3g15200 |
| *BVRB_1g010630* | 2-oxoisovalerate dehydrogenase subunit alpha 1, mitochondrial |
| *BVRB_1g011310* | Serine/threonine-protein kinase STY46 |
| *BVRB_1g012310* | Probable LRR receptor-like serine/threonine-protein kinase At1g34110 |
| *BVRB_1g012490* | Imidazole glycerol phosphate synthase hisHF |
| *BVRB_1g015750* | Isocitrate dehydrogenase [NADP] |
| *BVRB_1g016200* | ATP synthase subunit alpha |
| *BVRB_1g016350* | BRISC and BRCA1-A complex member 2 |
| *BVRB_1g020200* | DNA-directed RNA polymerases II, IV and V subunit 6A |
| *BVRB_1g020420* | Leucine--tRNA ligase, cytoplasmic |
| *BVRB_1g020800* | BTB/POZ domain-containing protein At2g30600 |
| *BVRB_1g022260* | Serine/arginine-rich splicing factor SC35 |
| *BVRB_1g022270* | O-fucosyltransferase 20 |
| *BVRB_1g022330* | Clathrin heavy chain 2 |
| *BVRB_1g023250* | Glutamate decarboxylase |

## Discussion

### CircRNA sequencing explains the mechanisms of drought resistance in sugar beet

The non-coding RNA sequencing offers a profound understanding of the anti-drought mechanisms in plants. Being crucial non-coding RNAs, circRNAs were previously regarded as abnormal splicing by-products (Cocquerelle *et al*., 1993; Jens, 2014). The vital functions of circRNAs in various biological events have been reported lately (Chen *et al*., 2017; Yin *et al*., 2017), with a few circRNAs capable of being translated into proteins or polypeptides (Legnini *et al*., 2017). The presence of circRNAs has been detected in mammalian cells (Salzman *et al*., 2012) as well as in the plants like arabidopsis, rice, soybean, and wheat (Chen *et al*., 2017; Wang *et al*., 2017; Yuan *et al*., 2018; Zhao *et al*., 2017). Nevertheless, there has been no complete elucidation of drought-responsive circRNAs in sugar beet so far. In the present study, the leaf morphogenesis circRNAs of sugar beet were identified in response to the drought challenge. They were characterized in a genome-wide manner, yielding a total of 563 circRNAs. The outcome corroborated prior works on soybean (776 of the entire 5,372 circRNAs were recognized from the leaves) (Zhao *et al*., 2017) as well as those on arabidopsis and rice (Lu *et al*., 2015; Ye *et al*., 2015). 432 of 563 circRNAs (76.7%) identified herein were of exonic type (Fig. 2A).

The correlation between circRNAs and stress response has been confirmed lately. For example, 62 wheat circRNAs, exhibited a specific pattern of expression upon dehydration challenge (Wang *et al*., 2017). Besides, the chilling injury response of 163 tomato circRNAs has been demonstrated, indicating a probable regulatory function of plant circRNAs in response to the cold challenge (Zuo *et al*., 2016). In the present work, there were eight upregulated, and four downregulated circRNAs (Fig. 4) among the 12 randomly picked circRNAs, showing agreement with the stress-specific patterns of expression for most of the plant circRNAs (Gao *et al*., 2015). Suggestively, the sugar beet circRNAs were probably implicated in the responses to drought stress.

As revealed by the qRT-PCR results, the expression trends of *novel_circ_0000591*, *novel_circ_0000695, novel_circ_0000764*, and *novel_circ_0000736* in sugar beet were linked positively to their respective parental genes (Fig. 4). This was in agreement with the previous findings (Ye *et al*., 2015). In the present study, the functionality of the parental genes was investigated by the KEGG and GO assessments under drought challenges. Multiple functions were observed for plenty of anticipated parental genes of circRNAs. According to the outcomes of GO analysis, the responses of sugar beet circRNAs toward drought challenges were linked to diverse functionalities involving various biological events, cellular components, as well as molecular functions (Fig. 5A). The KEGG assessment revealed 13 pathways that were linked to the drought tolerance of sugar beet. For instance, in the course of acclimation to drought sucrose, photosynthesis and synthesis become less efficient as well as N and C metabolisms exert crucial effects through interplays with other photosynthetic processes (Yang *et al*., 2019). Respiration in plants is influenced by drought, and prolonged periods of drought probably lead to the uncoupling of oxidative phosphorylation (Liu *et al*., 2020a). Both the KEGG and GO outcomes suggested the probable regulatory role of circRNAs in the sugar beet response to drought.

### CeRNA networks could provide new insights into the regulatory roles of ncRNAs during drought acclimation

CircRNAs were capable of sponging miRNAs and controlling the gene expression through their target mRNA regulation (Hansen *et al*., 2013; Wang *et al*., 2021b). For example, encompassing 16 presumed binding sites for miR138 by Sry circRNA and repression of the miR7 activity by ciRS-7/CDR1 suggested the miRNA sponging activity of circRNAs (Hansen *et al*., 2013). There are 1,861 circRNAs in plants that could probably sponge the miRNAs (Chu *et al*., 2017). In soybean, the speculated binding sites for 92 miRNAs were encompassed by 2,134 circRNAs (Zhao *et al*., 2017). Cleavage mediated by miRNAs has been suggested as a possible contributor to the low abundance and decomposition of plant circRNAs (Li *et al*., 2017a). To date, circRNAs have usually been considered as the ceRNA (competing endogenous RNA) molecules for predictive analyses (Liu *et al*., 2017; Wang *et al*., 2017). As displayed in Supplementary Table S3, presumed binding sites for 166 miRNAs were discovered for 197 circRNAs in the present study. Fifty-five of these circRNAs possessed over one such binding site, resembling the prior finding on *Brassica rapa* (Liu *et al*., 2022a). According to Wang et al. (2020), best efficacy could be occasionally sustained by the mismatches in miRNA target sites rather than the ideally paired sites.

Besides, for certain targets, natural selection either tolerate or has maybe opted for the suboptimal efficacies of miRNAs (Liu *et al*., 2014). The efficacies of several speculated miRNA target sites in the circRNAs thus appeared to be weak for the target cleavage in the present study, despite their ability to serve as the miRNA mimics. IPT, one of the predicted target mRNAs, was capable of enhancing the cold resistance in sugar cane (Belintani *et al*., 2012) and anti-drought traits of peanut (Qin *et al*., 2011). Besides, in soybean, the LT and drought responses are attended by the transcriptional factors DREB and MYB (Kidokoro *et al*., 2015; Su *et al*., 2014). Presumably, the circRNAs exert certain functions when encountered with drought and other environmental stimuli.

Recently, the regulation of circRNAs, miRNAs, and mRNAs has been confirmed in the case of various diseases (Lin and Yuhan, 2018). In the initial research concerning the SSR identification within the infection-responsive lncRNAs, the lncRNAs were definitively found to be useful in the breeding of Brassica crops (Summanwar *et al*., 2020). Nevertheless, there has been no extensive establishment of the circRNA–miRNA–miRNA regulatory meshwork in the plants. To understand the circRNAs functionalities and the ceRNA network in sugar beet under drought stress more clearly, a putative meshwork of circRNA-miRNA-mRNA was created. Here *novel_circ_0000443* and novel_circ_0000442 were chosen with their corresponding miRNA *ath-miR157d* and target genes of *ath-miR157d*. These stress-responsive target genes probably exerted a crucial function in tolerating drought. Some target genes of *ath-miR157d* were further examined after the statistical processing. Through targeting *ath-miR157d*, the *novel_circ_0000442* and *novel_circ_0000443* probably played a sponge role., This was how the control over the expression of its target genes was achieved. These included *BVRB_1 g004570, BVRB_1 g005450andBVRB_1 g005790*, whose functions were to code for U-box domain-containing protein 33, protein CLT2, and methionine adenosyltransferase 2 subunit beta, respectively. Further research is required to clarify the in-vivo associations among the *novel_circ_0000442* and *novel_circ_0000443*, *ath-miR157d*, and target genes, including *BVRB_1 g004570, BVRB_1 g005450* and *BVRB_1 g005790*. Moreover, future studies should focus on the downstream actions of these genes to better elucidate the anti-drought mechanism. Our findings offered novel ideas regarding the resistance mechanism of Chinese cabbage against the club root disease.

## Supplementary data

The following supplementary data are available at *JXB* online.

Table S1. Primer sequences used for qRT-PCR.

Table S2. Characterization of sugar beet circRNAs.

Table S3. MiRNA binding site of circRNAs.

## Acknowledgements

We acknowledge the time and expertise devoted to reviewing this manuscript by the reviewers and the members of the editorial board. We acknowledge some workers and staff members (Shanxi Agricultural University) for their assistance during our experiments. We thank Sagesci (www.sagesci.cn) for its linguistic assistance during the preparation of this manuscript.

## Author contributions

CLZ and CZ: conceptualization, design, writing with input from all authors; CLZ, ZG, SZ and JC: conducting the experiments; CLZ: data analysis. All authors read and approved the manuscript.

## Data availability

All data presented or analyzed in this article are available online through figshare and in the supplementary data. The raw sequencing data of sugar beet circRNAs have been submitted to the NCBI Gene Expression Omnibus (GEO accession numberGSE205327).

